# Detection of Unlabeled Micro- and Nanoplastics in Unstained Tissue with Optical Photothermal Infrared Spectroscopy

**DOI:** 10.1101/2024.11.11.622943

**Authors:** Kristina Duswald, Verena Pichler, Verena Kopatz, Tanja Limberger, Verena Karl, David Wimberger, Robert Zimmerleiter, Wolfgang Wadsak, Mike Hettich, Lukas Kenner, Markus Brandstetter

## Abstract

In this study, we investigate the efficacy of Optical Photothermal Infrared (O-PTIR) spectroscopy, also known as mid-infrared photothermal (MIP) microscopy, for the detection of micro- and nano plastics (MNPs) down to diameters of 250 nm in mammalian tissues. Experiments with both *in vitro* 3D cell cultures derived from HTC116 colorectal cancer cell line and *in vivo* mouse tissue models were conducted to evaluate the spatial resolution limits and quality of spectra that formed the basis for label-free and non-destructive identification of MNPs. Our findings demonstrate the superior resolution of O-PTIR in imaging individual particles of 250 nm in mouse kidney tissues, surpassing the capabilities of traditional FTIR spectroscopy, which was applied as a reference technique. Furthermore, we introduce a semi-automated image analysis that incorporates machine learning algorithms to accelerate the detection process, thus improving throughput and minimizing the potential for human error. The results confirm that O-PTIR produces high-quality, artefact-free spectral images in a contact-less manner and significantly outperforms FTIR in terms of spatial resolution and signal-to-noise ratio in complex biological matrices.

## INTRODUCTION

Microplastics are an increasing concern in daily life, absorbed by living organisms and found globally - on land, in the air, in bodies of water ^1–3^, and even in remote places such as Arctic areas ^4^, isolated regions of the Pyrenees Mountains ^5^, or Arctic beaches on remote islands.^6^ The full extent of the consequences remains largely unknown.^7,8^ A WHO ^9^ study recently summarized research on human exposure to micro- and nanoplastic (MNP) particles, highlighting the need for reliable detection methods for particles under 10 μm. Particles in that size range can easily be taken up, primarily via ingestion ^10,11^, but also through inhalation ^12^ or by breaching the cutaneous barrier ^13^, leading to tissue accumulation.

Recent studies suggest that frequent exposure to high levels of microplastics may increase the risk of diseases like colorectal cancer (CRC).^14,15^ Several studies have suggested that microplastics promote metastasis in CRC ^16^ and breast cancer cells.^17^ These particles can remain in the cells and may even be passed on during cell division. It has recently been shown that cholesterol molecules facilitate MNPs in breaching the blood-brain barrier in mice.^18^ Kopatz ^18^ and Kaushik ^19^ have highlighted the threat posed by MNPs, which can cross the blood-brain barrier and potentially cause severe neurotoxicity. By conducting labelfree determination of cancerous and non-cancerous tissues in human bladder surgical specimens using Raman spectroscopy, Krafft ^20^ demonstrated the utility of this technology for intraoperative tissue detection and simultaneously identified the presence of microplastics, pigments, and other foreign materials within the tissue samples. Since MNPs are abundantly found in human tissues, there is a pressing need to develop dedicated detection methods that accurately assess their presence and impact. However, current analytical techniques for MNP detection and their interactions with living organisms are still nascent.^8^ Thus, fast and reliable methods for localization and identification of MNPs in biological samples are prerequisites for better understanding their impact on our health. Various studies have suggested methods to address these questions, at least partially, such as pyrolysis-based techniques ^21^, microscopic techniques ^22,23^, and vibrational spectroscopy. Concerning the latter, Fourier-Transform Infrared Spectroscopy (FTIR) ^24,25^ and Raman spectroscopy ^26^ have become reliable tools for identifying microplastics based on their unique chemical fingerprint. However, each method has its respective strengths and weaknesses. FTIR is usually only applied to detect particles with sizes above 10 μm due to a limited spatial resolution imposed by the optical diffraction limit and can also suffer from artefacts in reflection measurements that are difficult to interpret. Conversely, Raman spectroscopy can detect particles below diameters of 10 μm unless the sample exhibits fluorescence, which can superpose the polymer characteristics in the spectrum.^26^ Attenuated Total Reflection (ATR)-FTIR might be a more robust alternative against artefacts and fluorescence than FTIR and Raman techniques. Still, the ATR crystal must be in contact with the sample surface.^27^ Therefore, this method has a high potential for causing cross-contamination and lacks the necessary spatial resolution. These issues have stimulated the development of a the Optical Photothermal Infrared Spectroscopy (O-PTIR) technique. The O-PTIR method combines IR spectroscopy principles with photothermal detection, where a sample absorbs a laser pulse, and the subsequent localized heating and accompanying expansion in the material is measured using visible light optics.^28^ This allows for the collection of IR-spectra beyond the IR-diffraction limit, thus enabling detailed chemical analysis at the submicron scale without needing contact with the sample.^29^

Research on microplastics detection in tissue has mainly focused on isolating polymer particles from biological material. To do this, the surrounding biological tissue is usually removed, making the particles more straightforward to analyze. Different chemicals can be used to dissolve the tissue surrounding the microplastics.^30–32^ However, this process is time-consuming and carries the risk of changing the chemical constitution of microplastics or even unintentional removal of particular polymer types, which might lead to false measurement results.^33^ Furthermore, this process results in the loss of crucial spatial information, such as the exact location of the microplastics within a tissue. Knowing where and in which cells microplastics accumulate in the tissue is crucial as it provides insights into how microplastics might affect the tissue or lead to changes within it and, consequently, to assess their potential impact on human health and disease.

This study demonstrates that MNPs with diameters as small as 250 nm can be precisely detected in mouse kidney tissue using the O-PTIR technology. This method allows for the detection of MNPs without isolating them, surpassing the current detection limits of similar techniques. This advancement closes the gap between high-resolution, time-intensive methods, such as electron microscopy, and faster optical spectroscopy techniques, thereby enhancing the applicability of reliable, position-sensitive MNP detection in tissues.

We present a comparative analysis of the absorption spectra obtained from FTIR and O-PTIR using polystyrene (PS) particles of defined and varied sizes. Subsequently, we demonstrate the detection of MNP particles within a controlled three-dimensional (3D) cell culture model (spheroids). This model helps to isolate potential influencing factors within a complex cell matrix akin to mammalian tissue. The cells are cultivated in a controlled environment, avoiding the generation of different functional tissues. This methodology is supported by the initial steps towards semi-automated data evaluation using a region-growing image segmentation algorithm. Finally, we illustrate the feasibility of detecting MNP particles in actual mouse tissue. We emphasize how the high lateral resolution, surpassing the diffraction limit for IR wavelengths, enables the detection of MNP particles in a spheroid and actual mouse tissue down to a size of 250 nm in diameter. This highlights the method’s potential for detailed environmental and biological analyses with straightforward and rapid two-wavelength scans.

## MATERIALS AND METHODS

### Micro- and Nanoplastic Particles

Commercially available unlabelled and labelled PS particles with a diameter of 10.39 ± 0.13 μm (spherical, aqueous suspension, 5% w/v, blue coloured), 1.14 ± 0.03 μm (spherical, aqueous suspension at 2.5% w/v, ex/em= 530/607 nm), 1.16 ± 0.04 μm (spherical, aqueous suspension at 5% w/v, red coloured), 0.24 ± 0.01 μm (spherical, aqueous suspension at 2.5% w/v, ex/em = 502/518 nm) and 0.200 ± 0.007 (spherical, aqueous suspension, 5% w/v, unlabelled) were obtained from Microparticles GmbH (Berlin, Germany). As published, all particles were characterized by measuring zetapotential, size, and polydispersity index (PDI) on a Zetasizer Pro device (Malvern Pananalytical, Malvern, United Kingdom).^16^ We will refer to the particle sizes in the following as 10 μm and 1 μm and 250 nm and 200 nm.

### HCT116 Spheroids

Human colorectal cancer cell line HCT116 (DSMZ, Braunschweig, Germany) was cultivated in a fully supplemented Minimum Essential Medium (MEM) culture medium (10% FBS, 1% P/S, 1% L-glutamine) at 37°C and 5% CO_2_ in a humidified atmosphere. For spheroid formation, HCT116 cells were suspended with the respective MNPs (MNPs as described before, final MNP concentration = 1 μg mL^-1^) in fully supplemented MEM media and seeded on Petri dish lids using the hanging drop method ^34^ at concentrations of 3 × 10^3^ cells per drop. The MNP-cell suspension was required to ensure equal distribution of the MNPs within the spheroid. Other methods, like ultra-low attachment plates or directly using agarose-coated plates, were unsuccessful, as the direct contact of the MNPs to the plastic surface of the cell culture plate led to uneven distribution within the spheroid. After 24 h, the medium was replaced by 10 μl of fresh media, and spheroids were transferred to agarose-coated 96-well plates with 100 μL of fresh media per well. MNP-treated spheroids (MNP concentration = 1 μg mL^-1^) were embedded in Tissue-Tek O.C.T. Compound (Sakura, Torrance, CA) on day 7, and 7.5 μm thick cryosections were prepared on Superfrost™ microscope slides (Epredia, Kalamazoo, US) using a Leica CM3050 S cryomicrotome (Leica, Wetzlar, Germany) and stored at -80°C. The samples were fixed two times with 4% paraformaldehyde (PFA) in phosphate-buffered saline (PBS) for 15 min and washed with PBS three times; afterwards, the cells were dehydrated by an ethanol row applying 50%, 70%, 80%, 95% and 3x 100%, each for 10 min.

### Preparation of *ex vivo* Samples

Kidneys from C57Bl/6J wildtype mice were extracted and post-mortem injected with 50 μL of PS-particle mix (0.3% w/v 10 μm blue colored particles, 0.03% w/v 1 μm red-coloured particles and 0.015% w/v 200 nm unlabelled particles in PBS). The organs were then fixed for 24 h with 4.5% PFA before the samples were further desiccated and paraffin-embedded with a modified isopropanol protocol.^35^ Sections of 5 μm were cut and mounted on regular glass slides (SuperfrostTM Plus Adhesion Microscope Slides; Epredia, Breda Netherlands). For removal of paraffin before analysis, samples were alternately heated up to 65°C and quickly dipped into 100% isopropanol.^35^ The process was repeated several times until the tissue sections appeared uniformly whitish, and the surrounding paraffin was removed.

### Fourier-Transform Infrared (FTIR) Spectroscopy

FTIR is the gold standard in infrared spectroscopy to investigate the molecular composition and structure (“chemical fingerprint”) of the sample of interest. Each infrared absorption band corresponds to characteristic molecular vibrations, thus giving both qualitative and quantitative information about the specific functional chemical group.

Reference spectra were obtained with a Bruker LUMOS I FT-IR microscope (Bruker, Billerica, Massachusetts, US), a fully automated IR device with a high-sensitivity, liquid nitrogen-cooled MCT detector. It enables transmission, reflection, and ATR mode, whereas the latter two have been used in this work. The microscope features an 8x objective, digital zoom up to 32x, a CCD camera, and a motorized stage for precise operation. The measurement spot size was controlled by adjusting the aperture. For PS-bead analysis, a 10 × 10 μm aperture was used, and a 30 × 30 μm aperture was chosen for imaging to ensure sufficient signal-to-noise ratio (SNR).

The ATR system uses a motorized single-bounce ATR germanium crystal, which is lowered onto the sample to interact with the sample surface. The infrared light reflects at the interface, creating an evanescent field that extends a few micrometers into the sample. This method provides high-quality, robust and undistorted spectra, making it ideal for determining IR absorption bands. To maximize the signal-to-noise ratio (SNR), a 100 × 100 μm aperture was used for ATR measurements.

All spectra from the FTIR-microscope were obtained by conducting 32 scans for both the background and sample measurements. These were averaged to calculate the final IR absorption spectrum using Beer-Lambert’s law, 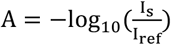, where I_s_ is a single beam reflection spectrum from the sample (particles) and I_ref_ is a reference or background single beam spectrum of a non-absorbing material (e.g. air or gold mirror). The spectral range of all spectra obtained with the FTIR covers the wavenumbers between 4000 cm^-1^ and 600 cm^-1^.

### Optical Photothermal Infrared Spectroscopy

The mIRage system (Photothermal Spectroscopy Corp., Santa Barbara, CA, US) was used to acquire Optical Photothermal Infrared (O-PTIR) spectra through a colinear pump-probe approach. A tunable pulsed EC-QCL laser, operating in the IR region from 3001 cm^−1^ to 2677 cm^−1^ and 1853 cm^−1^ to 933 cm^−1^, causes localized heating and thermal expansion in the sample. This photothermal effect is detected by a 532 nm CW-laser, with signal amplitude proportional to absorbed IR power, enabling quantitative absorption spectra. A lock-in detection modulation scheme improves SNR.

Typical optical spot sizes are 0.4 μm FWHM for the detection laser and 6.5–12 μm for the IR laser. O-PTIR spectra shown are averages of three scans. While Raman spectroscopy could be done on the same spot, further experiments were abandoned due to long integration times and fluorescence issues.

### Image Processing

The applied region-growing algorithm ^36,37^ is a segmentation technique that starts from seed points and expands to neighboring pixels based on criteria like intensity similarity or connectivity. This method includes adjacent pixels with similar properties to the seed point, forming a region until no more pixels meet the criteria. Post-processing techniques, such as morphological opening and closing, refine the segmentation by removing small artifacts and smoothing region boundaries. Opening, involving erosion followed by dilation, removes small noise and disconnects narrow connections, while closing, a dilation followed by erosion, fills small holes and smooths edges^38^. A rectangular kernel defines the neighborhood structure and guides morphological transformations for refinement, determining the scale and orientation of features affected during the process.

Hyperspectral imaging is also used to analyze small particles, evaluated using spectral descriptors - features derived from the spectrum of an image or signal, typically obtained through transformations like the Fourier or wavelet transform, capturing frequency or scale information. By analyzing spectral signatures at each pixel, these descriptors provide detailed information on the chemical composition and physical properties of particles, helping to analyze structure, patterns, texture, shape recognition, and signal processing. ^24^

## RESULTS AND DISCUSSION

Initially, we compare the spectral absorption characteristics of FTIR and O-PTIR for the MNP beads. Then we discuss experiments on MNPs incorporated into spheroids, which increased the sample complexity while providing a well-defined and controlled environment. Finally, we examine MNPs in tissue from mouse kidneys to demonstrate the method’s capabilities *in vivo*.

The experiments were performed with four types of polystyrene (PS) beads of spherical shape, with average diameters of 10 μm (blue label), 1 μm (fluorescence or red label), 250 (fluorescence label), and 200 nm (unlabeled). Details on the MNPs and sample preparation are described in the methods section.

### FTIR and O-PTIR performance on PS-Beads

In this section, O-PTIR results are compared to measurements in reflection and ATR measurement geometry using a conventional FTIR microscope. A schematic of the size relations between FTIR (red) and O-PTIR (green) detection spots and the measured MNP particles is shown in Figure 1 a), which also indicates the improved spatial resolution of the O-PTIR method. Although in the O-PTIR system, the spot size limit of the pump beam is fundamentally the same as in the classical FTIR microscope, the spot size of the laser beam used for detection is much smaller (∼500 nm). The corresponding FTIR reflection (green line), and O-PTIR (red line) spectra for every particle size are presented in Figure 1 b) and labelled accordingly. As a reference, the IR absorption spectrum taken in ATR measurement geometry is shown as a blue line at the top. It shall be noted that ATR spectra exhibit a slight wavelength dependent spectral shift of band positions, a well-known effect in IR spectroscopy. For additional characterization, we also applied the Raman spectroscopy method for MNPs; however, strong auto-fluorescence in the mouse tissue prevented any meaningful results. All shown spectra are averaged from three subsequent measurements, with measurement times of 60 s (FTIR reflection), 60 s (FTIR-ATR), and 12 s (O-PTIR). It is clearly visible that FTIR measurements taken in reflection geometry are prone to artefacts, mainly caused by optical scattering effects ^39^, baseline and band shifts ^40^, and particle geometry, while the spectra obtained with O-PTIR are significantly less affected. The resulting spectra show a clear chemical imprint of the PS beads with a superior signal-to-noise ratio (SNR) even at a shorter measurement time of the O-PTIR system. To corroborate the obtained spectral information, a test measurement with an FTIR-ATR was conducted on the beads with an average diameter of 10 μm. The aperture size for all FTIR reflection and FTIR-ATR measurements was 10 × 10 μm, which matches the size of the 10 μm particles and is the smallest aperture that results in a viable SNR.

**Figure 1:**
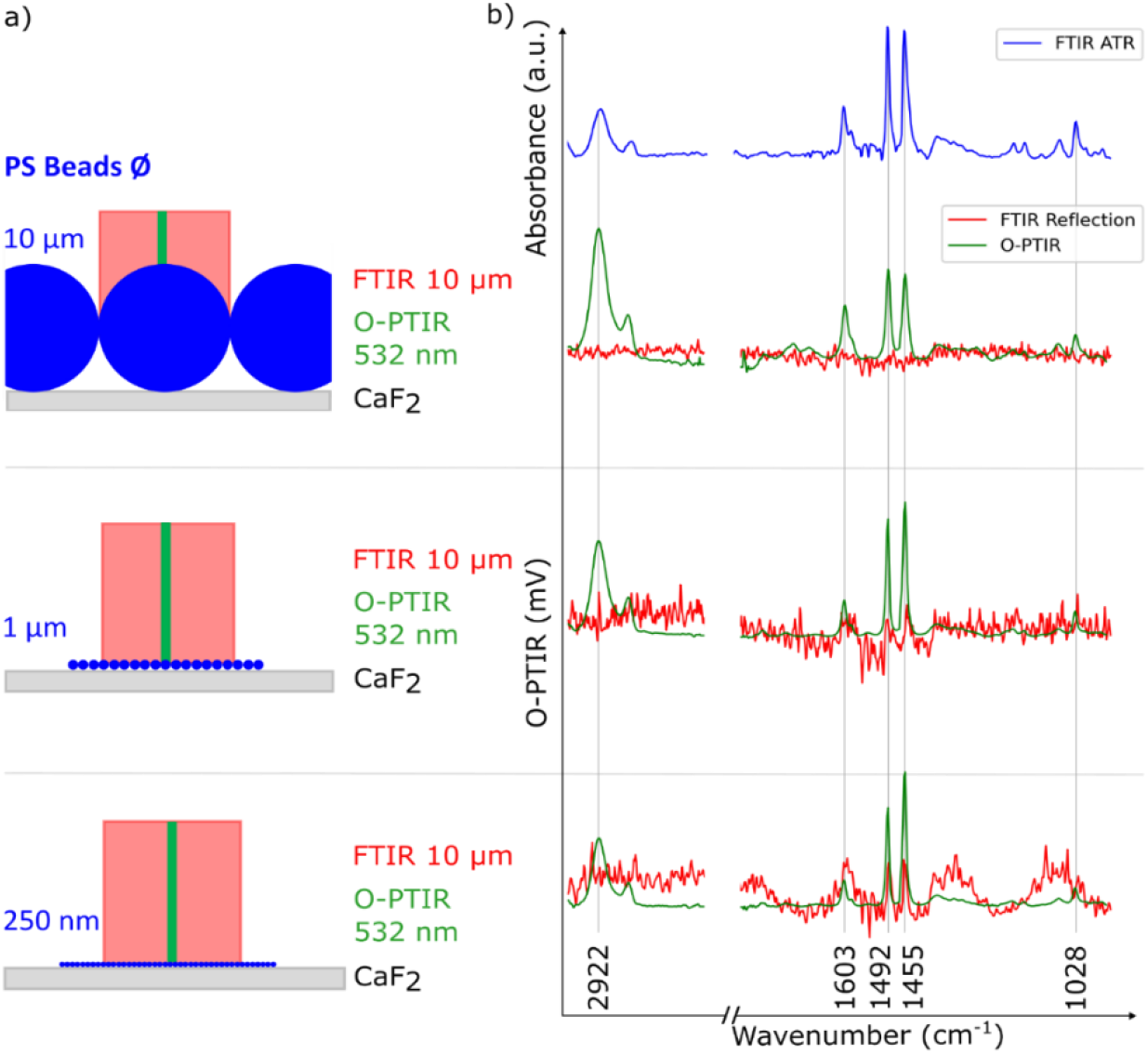
Comparison of the IR absorption spectra taken with three different methods – FTIR reflection (red), O-PTIR (green) and a reference FTIR-ATR (blue) for three different PS particle sizes (10 μm, 1 μm, 250 nm). In a) we show a schematic illustration of the field of view used: FTIR (red) and O-PTIR (green). Part b) shows the FTIR-ATR reference spectra taken from 10 μm beads followed by the comparison of FTIR reflection and O-PTIR. The top two spectra were taken from 10 μm particles, the middle from 1 μm particles, and the bottom from 250 nm particles.

A comparison of the obtained spectra shows an excellent agreement between the O-PTIR and FTIR-ATR results. In contrast, FTIR reflection spectra barely contain usable information due to an insufficient SNR. Using thermal light sources for classical mid-IR microspectroscopy requires apertures that reduce the SNR creating a trade-off between spatial resolution, SNR, field of view, and acquisition time, which is particularly problematic in low brightness scenarios like the mid-IR fingerprint region or in back-reflectance measurements such as *in vivo* diagnostics.^41^ Additional influence factors for this observation are the Rayleigh and Mie-scattering regimes for smaller and larger particles and the geometric scattering effects of the probe pulse at the used wavelengths. A thorough quantitative analysis of these effects, requiring detailed modelling of the detection process is outside the scope of this work and has been conducted elsewhere.^39^ Although the O-PTIR and the FTIR-ATR methods are comparable in terms of their spectral shape and SNR, the ATR method inherently demands that the crystal is in contact with the sample, which introduces the risk of cross-contamination and could cause alterations in the investigated samples due to the mechanical contact. FTIR-ATR also lacks the necessary spatial resolution unless combined with other techniques. Therefore, we used FTIR-ATR only as a complementary method for control measurements in the context of MNP detection.

It is clear from the data shown in Figure 1 b), that O-PTIR measurements do not exhibit the typical artefacts known from FTIR reflection spectra, which allows more accurate and consistent measurements across various sizes and shapes of MNPs. Furthermore, the O-PTIR approach allows a significantly higher spatial resolution, which is especially useful for the detection of MNPs.

### MNP (PS-Beads) in HCT116 spheroids

As a next step, based on the results presented in the previous section, MNPs with 1 μm and 250 nm diameters were incorporated within spheroids and measured using O-PTIR spectroscopy. This approach provided the basis for disentangling the cells and MNPs spectral fingerprints before proceeding to MNPs in mammalian tissue. Figure 2 a) compares an image from optical white-light microscopy and O-PTIR measurements. The white rectangle marked regions give a magnified view of the samples (Figure 2 b). The colour bar indicates the ratio between the absorption amplitudes at the selected spectral bands located at 1455 cm^-1^ and 1660 cm^-1^. The band at 1660 cm^-1^ corresponds to the Amide I band, primarily resulting from the C=O stretching vibration of peptide bonds in a biological sample - the band at 1455 cm^-1^ results from methylene vibrations from the PS beads. The spots that appear in red colour indicate the presence of one or several PS beads (white arrows). Figure 2 c) compares the spectra obtained on individual beads, pristine spheroids, and polystyrene (PS) MNPs embedded in a spheroid and indicates the positions of the wavenumbers that were selected for the ratio images. The blue spectrum in Figure 2 c) shows the characteristic absorption bands of the spheroid, while the grey spectrum was taken on individual PS beads. The red spectrum represents PS measured within a spheroid, showing a mixed spectrum from both entities. Although the presence of the spheroid created a background signal in the acquired IR spectrum, the absorption bands of the PS beads could still be distinguished unambiguously. As evident from the acquired spectral data, the ratio between the respective absorption peak amplitudes located at wavenumbers of 1455 cm^-1^ and 1660 cm^-1^ serves as a highly selective indicator for the presence of MNPs in spatial scans and the resulting images. Although a full spectrum was acquired at each pixel, reducing the scanning process to two distinct spectral features significantly improved the imaging speed. It took about 120 minutes to measure the two images of the relevant wavenumbers which resulted in a full image of 1883 × 1519 pixels and had an exact size of 470.5 μm width and 379.5 μm height. Built on this finding, further O-PTIR measurements were conducted on MNPs with diameters of 1 μm embedded in spheroid slices with a thickness of 10 μm, efficiently demonstrating the system imaging capabilities with chemical sensitivity.

**Figure 2:**
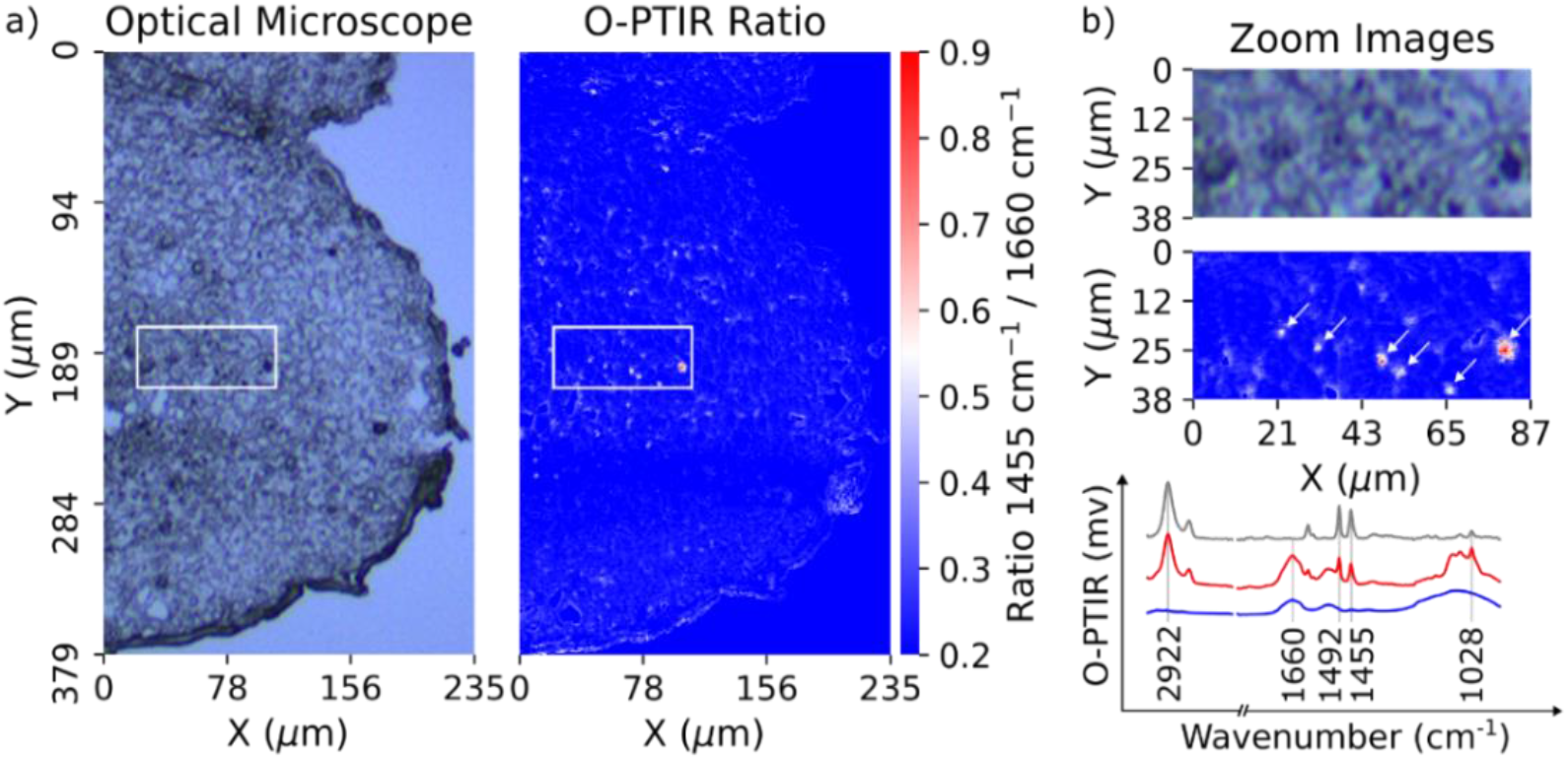
HCT116 spheroid was grown in fully supplemented MEM medium spiked with 1 μm PS spheres (1 μg mL^-1^). a) the left half shows a microscope image and the right half a false-colour image of the same area where the colour is given by the ratio of the absorbance peaks at 1455 cm^-1^ (characteristic for PS) and 1660 cm^-1^(biological tissue). B) The magnified marked areas show a microscope image (top) and a false-color image (bottom) indicating PS-MNPs. c) provides O-PTIR spectra from individual PS beads, embedded PS beads, and non-treated HCT116 spheroids. The grey spectrum is from a single PS bead, the blue is from a cell region without PS, and the red shows PS beads within the spheroid, clearly distinguishing both PS and cell features.

In the magnified view in Figure 2 b) the advantage of a chemically selective image of the embedded MNPs compared to the chemically non-selective visual image becomes apparent. A full spectrum was recorded for each localized polymer bead, and a random spot in the surrounding cells was used as negative control for validation. The optical microscopy image displays various structures and features, which can easily be mistaken for MNP contamination. However, in most cases, these are cellular structures or artefacts from sample preparation. In contrast, the chemical image focuses on the feature of interest (PS) and effectively ignores all the structural features of the cell. The false-coloured image was built solely on chemical information due to specific absorption features in the IR, clearly highlighting the areas where PS is present in the sample. The O-PTIR image (Figure 2) is based on pixel-wise information, assigning each pixel a continuous intensity value that represents the ratio described above.

We further aimed towards semi-automation through binary segmentation to accelerate the data analysis process. In this binary segmentation, Class 1 represents the particle of interest, such as PS, and Class 0 encompasses everything that does not match the value of the particle of interest. This method simplifies the evaluation by categorizing each pixel as either the particle of interest or not, enhancing the speed and accuracy of the analysis while integrating seamlessly with the transition towards semiautomated evaluations. Transitioning from manual to semi-automated evaluation processes improves efficiency and accuracy, allowing for rapid, consistent analysis while freeing up valuable human resources to focus on complex, judgment-based assessments. This shift not only streamlines routine evaluations but also ensures a more reliable and scalable approach to decision-making.

In Figure 3 we show the results of the segmentation process and their effects on particle detection. As mentioned above, the first image is based on the calculated absorption peak ratio we used as the initial point for feeding into the segmentation algorithm.

**Figure 3:**
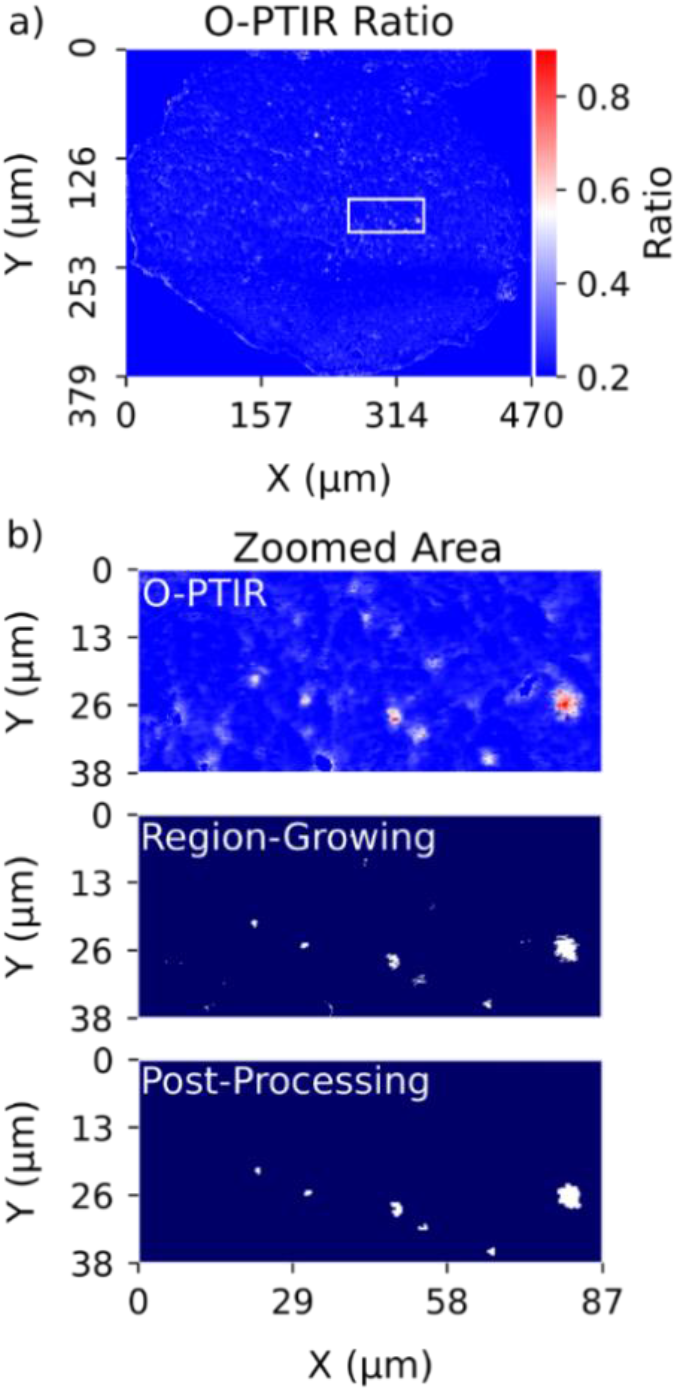
HCT116 spheroid grown in fully supplemented MEM medium with spiked 1 μm PS spheres (1 μg mL-1). In a) false-colour image indicating the ratio of absorbance at 1455 cm-1 and 1660 cm-1 is displayed. Red shows a high presence of the characteristic bands of PS. For clearer visibility b) display only zoomed-in area of the full image. The middle zoom shows the segmentation of the ratio image after applying the region-growing algorithm. This segmentation was then post-processed (bottom) to assign individual pixels.

We applied a region-growing algorithm, which started from a set of seed points and expanded outwards by appending nearby pixels with similar properties, as in our case ratio values, to form larger regions. This method iteratively groups pixels or subregions into regions based on predefined criteria, effectively segmenting the image into distinct areas. The results of the region-growing algorithm are shown in the magnified area of Figure 3 b). Finally, a post-processing step was carried out to further refine the results. Therefore, an opening was used to remove slight noise and artefacts to fill in small holes and gaps in the segmented regions. These steps used a rectangular kernel, whose size could be adjusted, to refine the segmentation, making it cleaner and more continuous (Figure 3 b), bottom). We tested the imaging capabilities of the O-PTIR system further in terms of achievable detection sizes of MNPs. 250 nm PS beads embedded in a spheroid were used to determine if the detection of individual beads was possible. We observed a tendency for aggregation of this 250 nm bead size in connection with Fetal Bovine Serum (FBS), which prevented us from imaging unclustered PS beads in the spheroid. In Figure 4 a), an agglomerate within a spheroid was examined in more detail using hyperspectral imaging (HSI). Here, we recorded a complete spectrum from each measured pixel in the wavenumber range from 3000 cm-1 to 2679 cm-1 and from 1857 cm-1 to 900 cm-1. Figure 4 a) also displays the spectral/chemical changes along the highlighted measurement positions. The characteristics of the typical PS bands (1455 cm-1) are visible in red areas, and in the blue regions, specific protein bands (1660 cm-1) are present. These relevant bands were used to obtain the ratio plot in Figure 4 b). As one moved further from the centre of the agglomerate towards the surrounding cells, the protein-specific bands became more pronounced while the PS-specific bands weakened (from 1 - 12). This fading effect indicates that the agglomerate of 250 nm PS beads is found deeper within the 10 μm tissue cross-section when moving away from the centre of the agglomerate.

**Figure 4:**
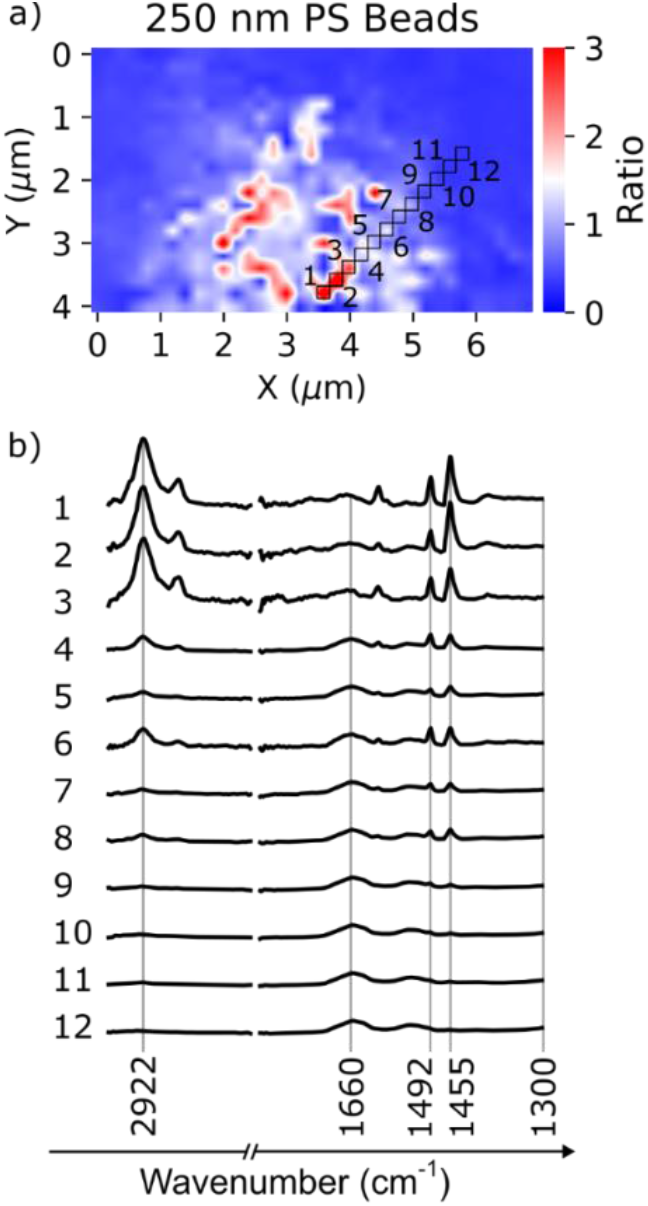
In image a) the ratio of bands 1455 cm^-1^/1660 cm^-1^ is depicted per pixel. It is a section of agglomerated 250 nm PS beads with a total diameter of approximately 4 μm. The black squares indicate measurement points along a line from the centre to the edge of the PS agglomeration. The spectra are displayed in b) and are numbered.

The findings on MNP-embedded spheroids show that chemically resolved imaging with O-PTIR is possible and can detect beads with diameters down to 250 nm. Since the measured beads were aggregated, only clusters of particles were detected. In the next section, we present further data to support our claim, where individual beads were imaged. It was essential for measurements on actual tissues that, although a background spectrum was present in the data, the PS-bead-specific absorption bands could be unambiguously identified inside the spheroid. Reducing the complete spectrum acquisition to specific bands is likely to further minimize imaging times.

### Detection of MNPs in Mouse Kidney Tissue

Building on the spheroid results, we moved to the next crucial step for actual biological and medical applications, i.e. investigations of *ex vivo* tissue samples. O-PTIR measurements were carried out on mouse kidney tissue embedded in paraffin.

Mouse kidney tissue was spiked post-mortem with PS beads with diameters of 10 μm, 1 μm, and 200 nm following the procedure described in the Methods section. Similar as for the spheroids, we compared the spectra of the different bead sizes in combination with the background of the mouse kidney tissue (Figure 5). Similar to the spheroids, the ratio between the 1455 cm^-1^ and the 1660 cm^-1^ band determined whether PS was presented throughout the kidney tissue. Consistent with our findings on the spheroid samples, the spectra of the beads with 1 μm diameter combine the spectral features of the surrounding biological material, here the mouse tissue, and the PS beads. Understandably, the contribution of the surrounding mouse tissue (5 μm thickness of tissue section) was more pronounced in the case of the smaller 200 nm beads. In the case of the 10 μm beads, however, the observed signal resembled a pure PS spectrum without any signatures of the surrounding tissue. This observation is consistent with the approximate size of the IR laser spot, i.e., for larger particles, the surrounding tissue was not excited and thus did not contribute significantly to the O-PTIR signal.

**Figure 5:**
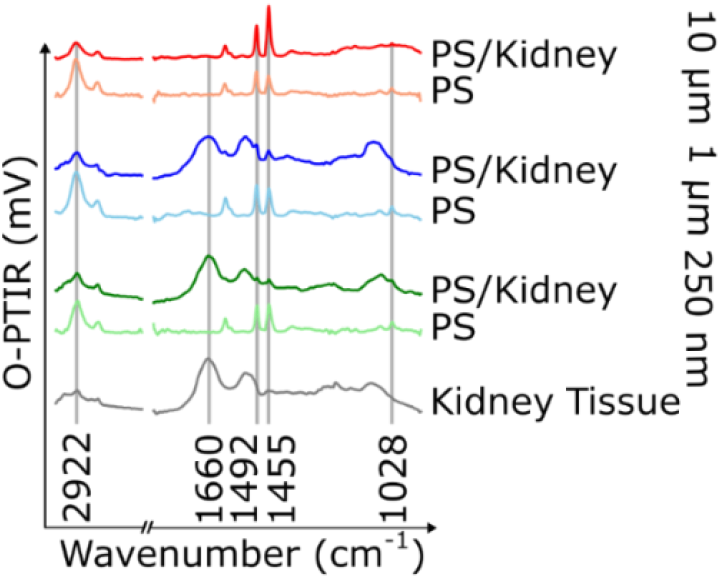
Comparison of O-PTIR signals for isolated spherical PS beads with diameters of 10 μm, 1 μm, and 200 nm with the signal of PS beads spiked into mouse kidney tissue. Vertical lines mark the characteristic absorption bands.

In line with our observations from the spheroid samples, the Amide I band at 1660 cm^−1^, depicted as a blue vertical line in Figure 5, was used as a reference for the biological material. Similarly, the characteristic absorption at 1455 cm^−1^ was leveraged to identify the presence of PS. Thus, the ratio of these distinct absorption bands, 1455 cm^−1^ and 1660 cm^−1^ again served to quantify the relative contributions of tissue and PS.

Figure 6 compares an optical microscope and an O-PTIR image false-colour image (embedded in Figure 6 a) based on the ratio at the two characteristic wavelengths for mouse tissue and PS. We also tested the same computational procedure (Figure 6 b) used in the spheroid segmentation process on the complex tissue of the mouse kidneys. After implementing the region-growing algorithm, significantly more noise remained in the kidney tissue compared to the spheroid sample. Our post-processing aimed to strike a balance where particles were properly segmented without losing too much information. While this allowed us to filter out a significant amount of noise, some background noise remained. It could even mislabel regions with PS particles in them as pristine tissue: the area where the 200 nm beads were localized was not recognized by the post-processing and was falsely labelled. This indicates that the ratio method worked for particles of 1 μm and 10 μm but did not provide enough information to unambiguously detect 200 nm particles in complex tissue.

**Figure 6:**
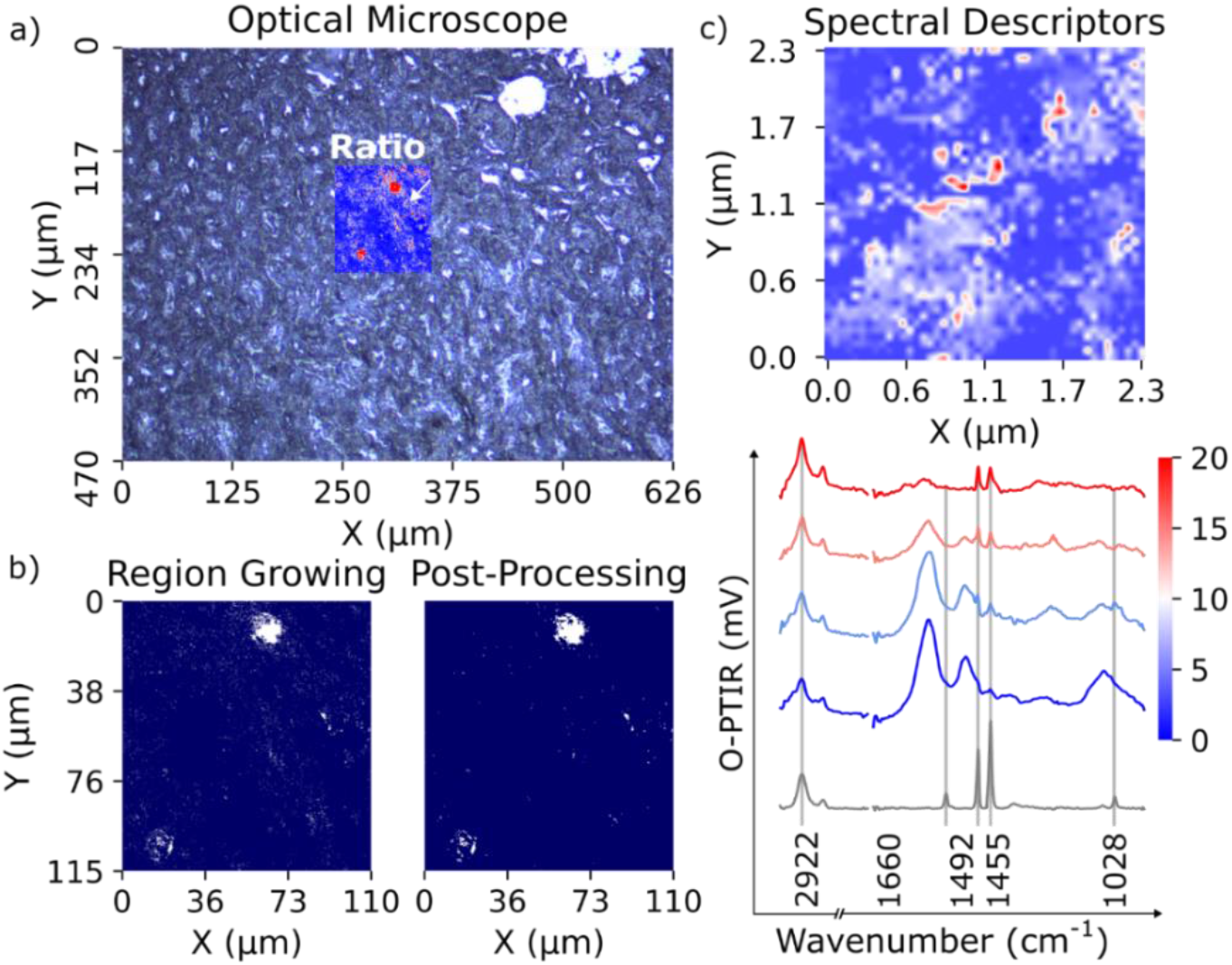
Mouse kidney tissue spiked with spherical 10 μm, 1 μm, and 200 nm PS beads. In a) the cross-section of a spiked mouse kidney is visible. The area of interest is overlapped with a ratio image (1455 cm^-1^/1660 cm^-1^). Growing segmentation and postprocessing are applied in the sub-images in b). The white arrow in the ratio image indicates the 2.3 × 2.3 μm area recorded as an HSI, which is illustrated in c). Therein, 200 nm particles in the mouse tissue become visible. Corresponding spectra confirming their chemical identity are shown in the bottom plot in c).

Based on this finding, we recorded a high-resolution hyperspectral image (Figure 6 c). For this map, we used two spectral descriptors ^24^ by using the software ImageLab (Epina GmbH): one at the wavenumber 1492 cm^-1^ and the other at 1455 cm^-1^.

The employed method calculates whether a triangle can fit under the respective absorption bands, thereby distinguishing unspecific spectral changes from those associated to actual IR bands The spectra in the graph correspond to the colour coding of the 2D map, indicating where the influence of PS is higher (red) or lower (blue). The results (Figure 6 c) demonstrate that PS particles of all sizes can be detected in mouse kidney tissue. Larger diameters were identified using the time-efficient ratio method as image contrast, while smaller particles with a 200 nm diameter required acquiring full spectral features for identification. A detailed analysis of the physical representation of the different particle sizes in the O-PTIR images is provided in the supplemental section (Figure S1). To emphasize the implemented technique’s impressive performance, we provide a comparison of the investigated region imaged by optical microscopy a), FTIR reflection b), and O-PTIR c) in Figure 7. The left picture displays the image captured with white-light microscopy in epi-configuration. Here, two beads with a diameter of 10 μm, indicated by white arrows are directly recognizable. However, this becomes more challenging with beads that are 1 μm in size, as indicated by the red arrow. Although the FTIR reflection measurement suggested the presence of MNPs all over the imaged area, the lack of spatial resolution prevented meaningful data extraction. The spectra of the red-appearing area were visually proofed, and there were no significant PS characteristics visible. An aperture size of 30 × 30 μm was chosen for the FTIR measurements, which was the smallest possible to obtain a usable signal in reflection mode. However, as previously mentioned, this led to significant artefacts in the absorption spectra. In stark contrast to the white-light microscopy and FTIR, the O-PTIR image shows a distinct contrast between the PS beads of 10 μm and 1 μm size and the surroundings, also clearly revealing the shape of the particles. The unmatched spatial resolution and measurement performance of the O-PTIR measurement method allowed a more in-depth analysis of the individual particles with different sizes present in the mouse tissue i.e. 10 μm, 1 μm, and 200 nm.

**Figure 7:**
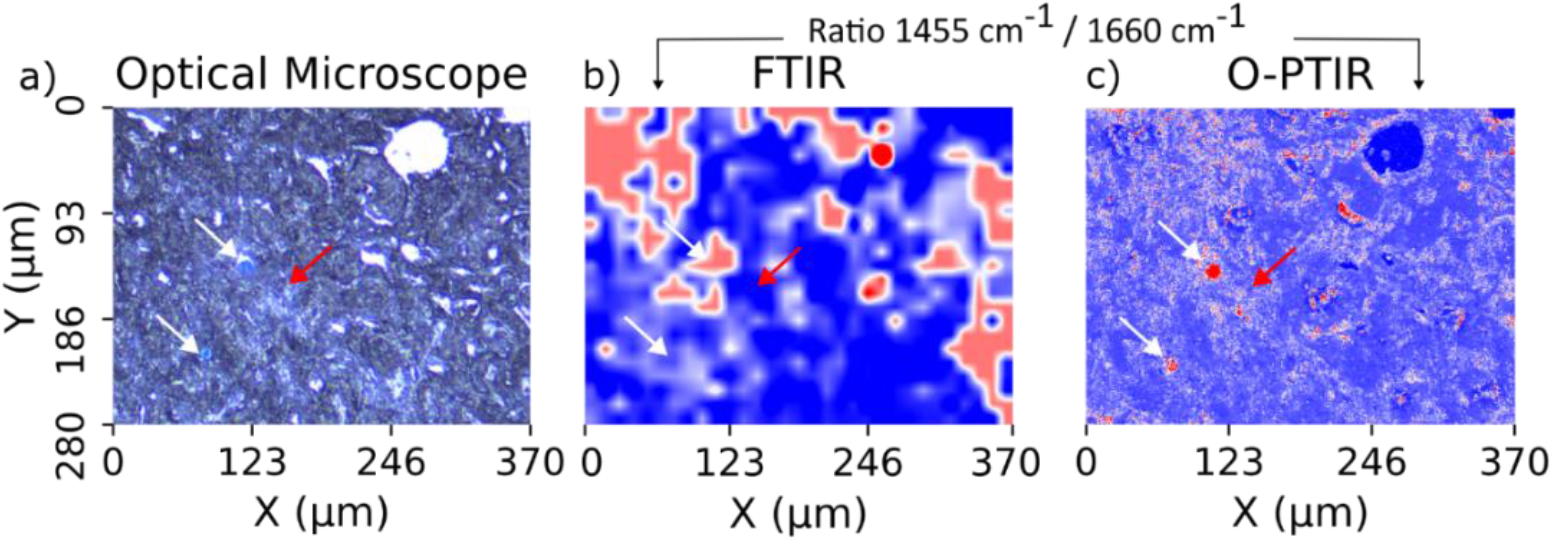
Comparison of imaging techniques in mouse kidney tissue section spiked with 1μm, 10μm and 200 nm PS-particles. The image a) shows a white-light microscopy image of the area, b) and c) show false-colour images obtained with FTIR- and O-PTIR microscopy, respectively. For the FTIR-image an aperture of 30 × 30 μm was used, while the O-PTIR image was taken with a spatial resolution of 500 nm.

## CONCLUSION

In this study, we employed O-PTIR spectroscopy to effectively identify microplastic particles (MNPs) with diameters of 10 μm, 1 μm, 250 nm and 200 nm – enabled by the significantly higher spatial resolution compared to FTIR microscopy. Artefacts, typical for infrared reflection measurements are not present in O-PTIR, which massively improves spectra interpretation. Unlike traditional FTIR, which requires spectral acquisition at each spatial point to build a hyperspectral image, O-PTIR can capture either full hyperspectral images or images at selected wavenumbers characteristic for a distinct polymer type. This flexibility, particularly the ability to acquire data at specific wavelengths, dramatically increases image acquisition speed, while still delivering essential chemical information.

Our results prove the capability of O-PTIR to detect microplastic particles in formalin-fixed paraffin-embedded (FFPE) tissue sections, a significant advancement over traditional methods that rely on tissue digestion and filtration. This allows for direct chemical imaging of particles within a tissue matrix, providing spatial information on their precise locations without the need for destructive sample processing. Remarkably, despite the inherent challenges in detecting particles as small as 200 nm, which are often difficult to identify even using traditional methods, O-PTIR proves capable, thereby presenting a viable alternative to FTIR.

This advancement offers valuable insights into the distribution of MNPs in FFPE tissue samples, providing histopathological context on particle accumulation and its potential implications. By offering enhanced spatial resolution and reduced artefacts, O-PTIR paves the way for more detailed and non-destructive analysis of microplastic contamination in biological tissues. Future work will focus on refining segmentation algorithms to improve detection speed and accuracy, further advancing our ability to monitor microplastic distribution in complex tissue samples.

## Supporting information

Individual plastic particles size profiles

## AUTHOR INFORMATION

### Corresponding Authors

* Markus Brandstetter

## Author Contributions

K.D. Methodology, Software, Validation, Formal analysis, Investigation, Data Curation, Writing – Original Draft, Visualization

V.P. Validation, Investigation, Resources, Writing – Review & Editing

V.Ko. Validation, Investigation, Resources, Writing – Review & Editing

T.L. Validation, Investigation, Resources, Writing – Review & Editing

V.Ka. Validation, Data Curation

D.W. Investigation, Data Curation, Writing – Original Draft, Review & Editing

R.Z. Writing – Review & Editing, Visualization

W.W. Conceptualization, Investigation, Project administration, Funding acquisition

M.H. Supervision, Writing – Original Draft, Writing - Review & Editing

L.K. Supervision, Writing – Review & Editing, Project administration, Funding acquisition

M.B. Conceptualization, Investigation, Supervision, Writing – Review & Editing, Project administration, Funding acquisition

## ACKNOWLEDGMENT

K.D., V.P., V.K., L.T., V.K., D.W., R.Z., W.W., M.H., L.K., M.B. acknowledges the support from MicroONE, a COMET Modul under the lead of CBmed GmbH, which is funded by the federal ministries BMK and BMDW, the provinces of Styria and Vienna, and managed by the Austrian Research Promotion Agency (FFG) within the COMET—Competence Centers for Excellent Technologies—program. K.D. and D.W. acknowledge support by research subsidies granted by the government of Upper Austria (Grant Nr. FTI 2022 (HIQUAMP): Wi-2021-303205/13-Au). Financial support was also received from the Austrian Federal Ministry of Science, Research and Economy, the National Foundation for Research, Technology and Development, and the Christian Doppler Research Association, as well as Siemens Healthineers for their financial and scientific support. L.K. was also supported by a European Union Horizon 2020 Marie Sklodowska-Curie Doctoral Network grants (ALKATRAS, n. 675712; FANTOM, n. P101072735 and eRaDicate, n. 101119427) as well as BM Fonds (n. 15142), the Margaretha Hehberger Stiftung (n. 15142), the Christian-Doppler Lab for Applied Metabolomics (CDL-AM), and the Austrian Science Fund (grants FWF: P26011, P29251, P 34781 as well as the International PhD Program in Translational Oncology IPPTO 59.doc.funds). Additionally, this research was funded by the Vienna Science and Technology Fund (WWTF), grant number LS19-018, and L.K. received support from a European Union Horizon 2020 Marie Sklodowska-Curie Innovative Training Network grant (ALKATRAS). L.K. is a member of the European Research Initiative for ALK-Related Malignancies (www.erialcl.net).

