## Supplementary material for "Detection of Unlabeled Micro- and Nanoplastics in Unstained Tissue with Optical Photothermal Infrared Spectroscopy": Individual plastic particles size profiles

### SUPPLEMENT MATERIAL

In Figure S1 false-colour images of selected areas for each particle size are shown. The color is determined by the ratio of the absorbance peaks at 1455 cm$^{-1}$ characteristic for PS and 1660 cm$^{-1}$ characteristic for biological tissue. Individual particles were recognized and confirmed by a complete spectrum acquisition. Special care was taken to locate isolated particles. The right-hand side shows the respective lateral mean intensity profiles in the x- and y-directions along the paths indicated in the false-colour image. We decided to use a mean spatial average for the profiles instead of a single-pixel profile to reduce noise. The image was recorded with a spacing of 100 µm, resulting in the averaging over each column or range being a factor of 10 greater than the length of the corresponding axis.

In Figure S1 a), the 10 µm particle is shown. Compared to spheroids, the increased complexity of the mouse kidney tissue is noticeable as an accumulation of noise around the 10 µm particle. A closer look at the obtained image reveals a particularity

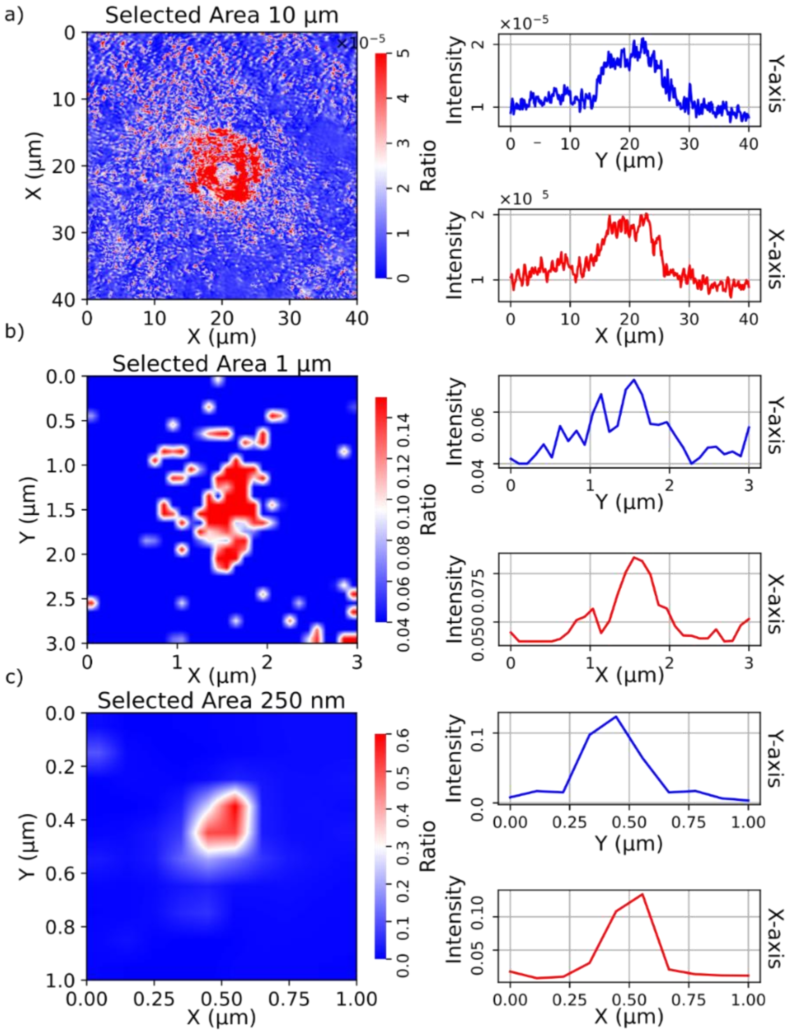


Figure S1: The left column shows the false-colour image of the different particle sizes (a) 10 µm, b) 1 µm, c) 200 nm) embedded in mouse tissue. The right column shows the particles' intensity profile. In addition to the height profiles, an averaged height profile along the x-axis and y-axis has also been created.

in detecting this bead size. The detected signal is smaller in the middle section of the particle, which results in a doughnut shape on the lateral profile (not visible in the averaged profile). We noticed a similar behavior for other particles of this size, which hinted towards an effect specific to the O-PTIR detection method related to the focusing optics and scattering effects of the IR laser beam from the curved particle geometry, due to the particle size comparable to the IR laser wavelength. However, closer investigation of this effect is outside of the scope of this article and will be further pursued separately. Despite this potential measurement artefact, a good agreement between the extracted particle size, measured as full width at half maximum (FWHM) of the averaged spatial profiles, and the expected particle size is achieved in the case where the spot size of the optical detection was considerably smaller than the particle diameter.

In Figure S1 b) a 1 µm particle is investigated. The extracted lateral profiles show a slightly smaller lateral size than expected. This could be due to a possible cut of the particle during the microtome processing or on the base of the surrounding and overlapping tissue, i.e. the measured cross-section of the particle could be decreased.

Figure S1 c) shows the O-PTIR image of an isolated 200 nm PS particle, the smallest particle size used in our experiments. This measurement method clearly demonstrates the excellent detection capabilities of small microplastic particles. The extracted profiles of the isolated bead are shown on the right-hand side, where we find a FWHM of 300 nm and 330 nm for the x- and y-direction, respectively. Since the spot size of our detection laser exceeded the particle size, we would have expected to find an image size of the particle with a dimension of at least the optical FWHM spot size of 400 nm. This assumes that the final image of the particle is a convolution between the particle size and the Gaussian optical beam. This would even be the case if we had assumed a point-like particle, i.e., a possible microtome cut of the particle could be ruled out as the origin of this result. The discrepancy shows that, like the observation mentioned earlier for larger particles, the details of the detection process require an improved understanding to deduce quantitative information when particles smaller than the optical spot size were investigated. We want to emphasize here that, although some uncertainties were introduced for small particles, different particle sizes can clearly be distinguished and therefore valuable statistical information on MNP-contaminated tissue can be provided.
